# A marine bacterium elicits the type III secretion-dependent nonhost hypersensitive response in a land plant

**DOI:** 10.1101/2023.06.20.545751

**Authors:** Boyoung Lee, Jeong-Im Lee, Soon-Kyeong Kwon, Choong-Min Ryu, Jihyun F. Kim

**Author notes:** Corresponding authors (J.F.K.); (C.-M.R.); (S.-K.K). These authors contributed equally to this work (Boyoung Lee, Jeong-Im Lee).

## Abstract

Active plant immune response involving programmed cell death called the hypersensitive response (HR) is elicited by microbial effectors delivered through the type III secretion system (T3SS). The marine bacterium *Hahella chejuensis* contains two T3SSs that are similar to those of animal pathogens, but elicited HR-like cell death in the land plant *Nicotiana benthamiana*, which was mediated by SGT1 and suppressed by AvrPto1. Thus, the land plant is capable of inducing the cell death in response to type III-secreted effectors of the marine bacterium it never have encountered, suggesting that plants may have evolved to cope with a potential threat posed by alien effectors.

**One-sentence summary:** A marine bacterium *Hahella chejuensis* harboring functional type III secretion systems induced nonhost hypersensitive response in a land plant *Nicotiana benthamiana*, which was mediated by SGT1 and suppressed by AvrPto1.

## INTRODUCTION

The most prominent strategy of plants to counteract pathogens is rapid and localized programmed cell death (PCD), termed the hypersensitive response (HR), which is similar to apoptosis in animal cells (Heath, 2000). HR initiates with recognition of a pathogen effector protein by a corresponding plant resistance protein with a nucleotide-binding leucine-rich repeat (NLR), in a process known as gene-for-gene resistance (Goodman, 1994). Plants also manifest the HR cell death as part of nonhost resistance against a broad spectrum of plant pathogens. Bacterial effectors are delivered into host cells through specialized type III secretion systems (T3SSs) (Kim, 2001). T3SSs are considered effective weapons for pathogenic bacteria to ensure successful infection by subverting the host immune system or by hijacking host metabolism (Grant et al., 2006; Xian et al., 2020). However, it is not known to science that any animal-pathogenic bacterium including *Yersinia* species is able to elicit HR in plants by injecting a T3SS effector into the cytoplasm.

To better understand the nonhost resistance of plants, we used *Nicotiana benthamiana* as a model plant and *H. chejuensis* as a model pathogen, for which the hosts do not include plants. We first assessed the expression of T3SS genes in *H. chejuensis* KCTC 2396 at different growth stages to explore the functionality of *H. chejuensis* T3SSs. We then infiltrated *H. chejuensis* cells into *N. benthamiana* leaves to observe HR-like necrosis. We further evaluated whether HR-like cell death elicited by *H. chejuensis* can be ameliorated when a bacterial effector that suppresses HR is expressed or a key component of R-gene-mediated resistance is silenced.

## RESULTS

### Overview of the T3SSs in the *H. chejuensis* Genome

The genome of *Hahella chejuensis* KCTC 2396^T^, a prodigiosin-producing member of oceanic *Gammaproteobacteria* (Lee et al., 2001; Kwon et al., 2010), contains two sets of genes encoding T3SS, which are highly similar to those in the virulence plasmids of animal-pathogenic *Yersinia* spp. (Figure 1A) (Snellings et al., 2001; Jeong et al., 2005). The export apparatus (SctRSTUV), inner and outer rings of the needle complex (SctJD/C), ATPase (SctN), the cytoplasmic sorting platform (SctKQ), the stator (SctL), the stalk (SctO), and even a controller of the release and translocation of Yop effector proteins (YopN) and its chaperone protein (SycN) (Cheng et al., 2001; He et al., 2004; Deng et al., 2017) were conserved in *H. chejuensis* (Figure 1A). To infer the phylogenetic affiliation of *H. chejuensis* T3SSs, we compiled all SctV sequences and analyzed their evolutionary relationships (Figure 1B and 1C). We designated these genes *hct* (*Hahella chejuensis* T3SS). As with YopN and SycN (data not shown), both HctVs formed sister branches to those of *Yersinia*-type SctVs (Figure 1C).

**Figure 1.**
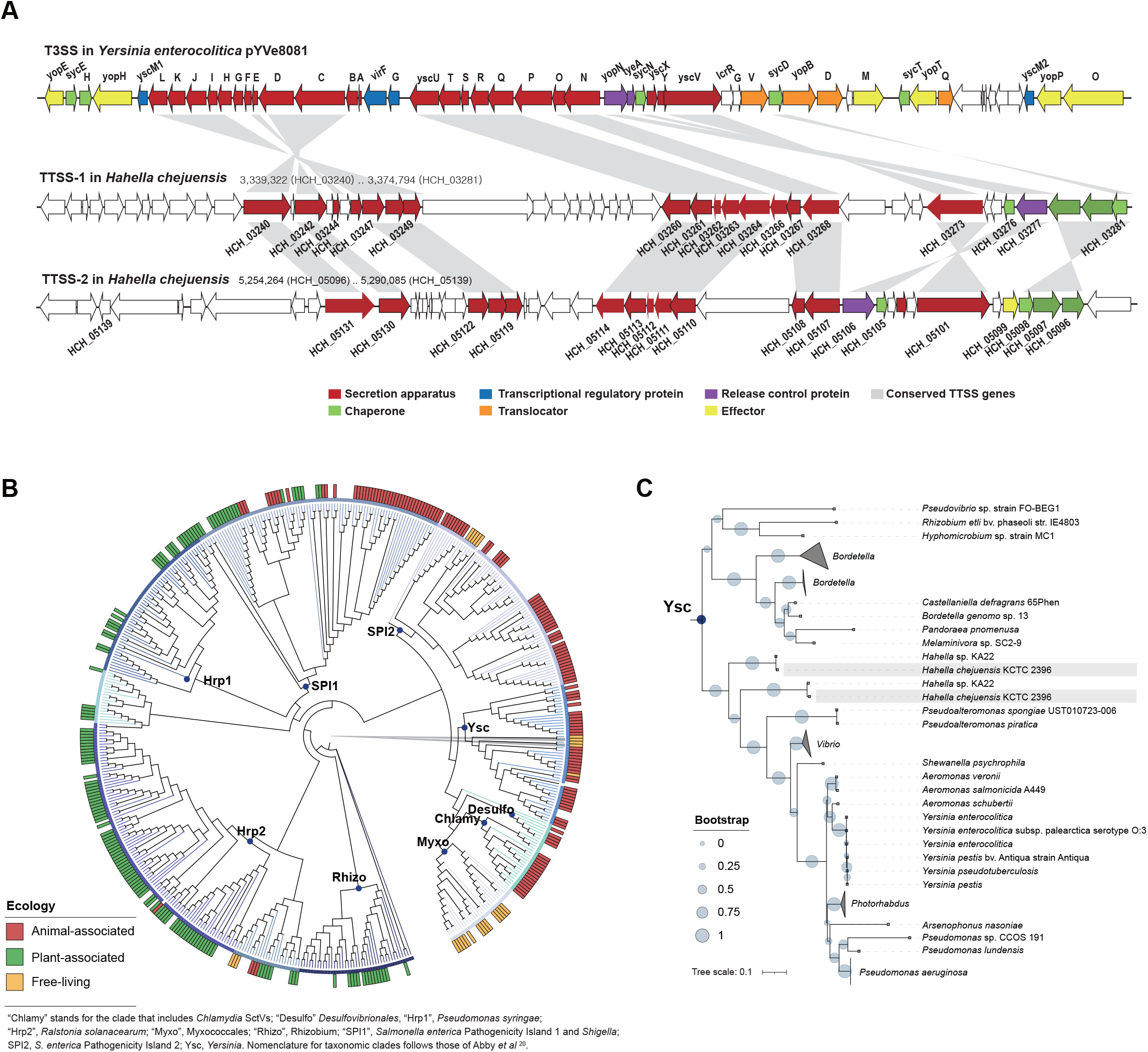
T3SSs in the *Hahella chejuensis* genome and the phylogeny of SctV. (A) Genetic organization of two T3SSs in the *H. chejuensis* genome with the archetypal T3SS of *Y. enterocolitica*. (B) A topological representation of a prokaryotic type III secretion inner membrane channel protein, SctV. The branches for *H. chejuensis* HctVs are in grey (Abby and Rocha, 2012). (C) Detailed maximum likelihood tree of the “Ysc” clade in (B).

### HR-like Cell Death in *N. benthamiana* Elicited by *H. chejuensis*

Considering that nonhost resistance of plants is a common form of disease resistance (Senthil-Kumar and Mysore, 2013) and the land plant *N. benthamiana* obviously is not a host for the T3SS-bearing marine bacterium *H. chejuensis*, we examined whether *N. benthamiana* would show HR against *H. chejuensis* by infiltrating the bacteria at different growth stages into the plant leaves. Surprisingly, necrosis appeared at 20 hours post inoculation (hpi), and the lesion became clear at 40 hpi only in areas where *H. chejuensis* in the late exponential phase (8 h) and in the stationary phase (10 h and 12 h) were infiltrated (Figure 2A and 2B). Infiltration of *Hahella*-extracted or synthetic prodigiosin at different concentrations resulted in no visible lesions (data not shown), ruling out that it is involved. In addition, *H. chejuensis* in the late stationary phase did not cause necrosis, even though the pigment production is higher. This HR-like cell death was comparable with the transcriptional patterns of *H. chejuensis* T3SS-1 genes, which showed bacterial growth dependency (Figure 2A, 2B and 2C).

**Figure 2.**
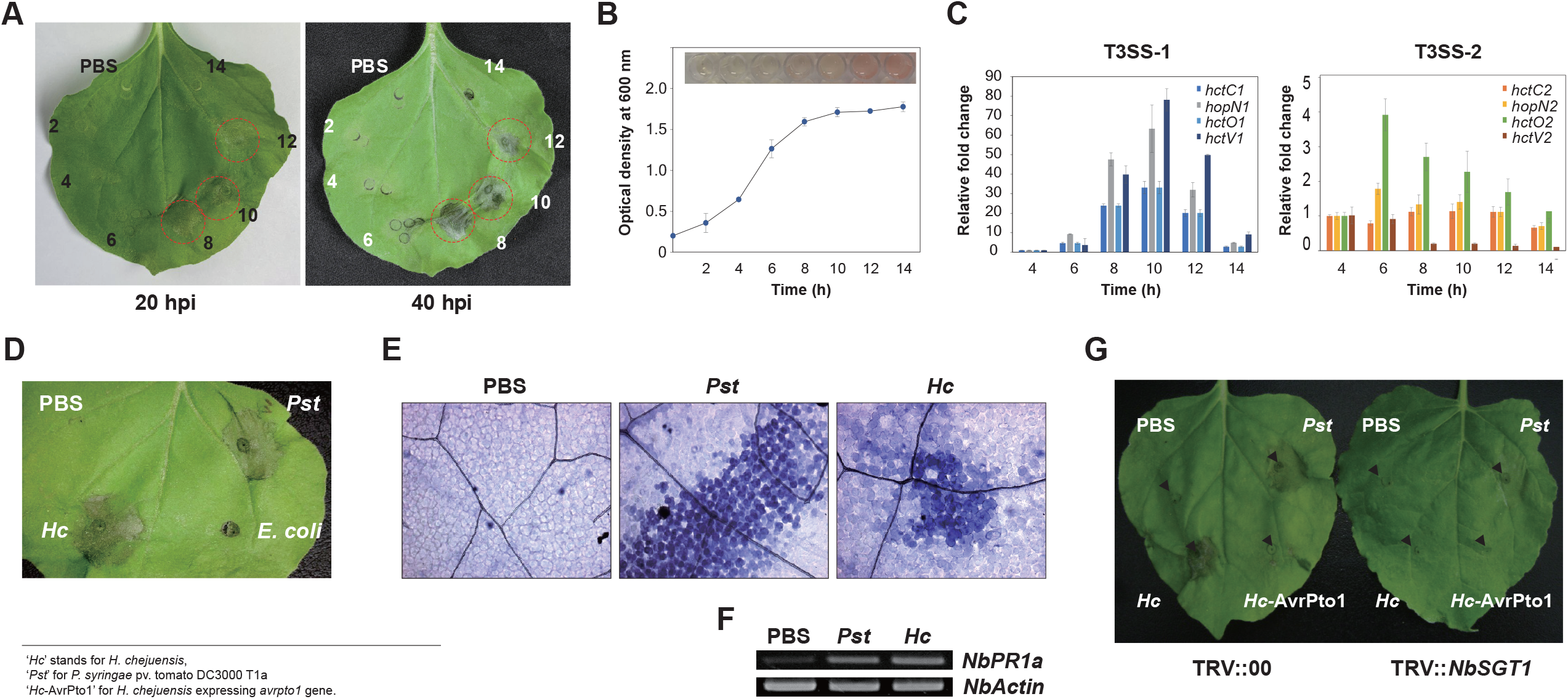
HR cell death in *N. benthamiana* elicited by the marine bacterium *H. chejuensis*. (A) HR-like cell death in *N. benthamiana* leaves at 20 and 40 hpi of *H. chejuensis*. (B) Growth patterns of *H. chejuensis* and red pigment production. (C) Expression of T3SSs genes in *H. chejuensis*. (D) HR-like cell death in *N. benthamiana* infiltrated with bacteria or PBS at 20 hpi. (E) Microscopic observation of the trypan blue-stained spots. (F) Expression of the *NbPR1a* gene at 12 hpi of bacteria. (G) Suppression of PCD by type III-secreted AvrPto1 or *NbSGT1* silencing in *N. benthamiana*.

We then tested whether the cell death in *N. benthamiana* was equivalent to PCD induced by a genuine plant pathogen by comparing the responses of *N. benthamiana* against *H. chejuensis, Pseudomonas syringae* pv. tomato DC3000 T1a (a tomato pathogen without *avrPto*), *Escherichia coli* EPI300 (a non-pathogen), or PBS buffer. Upon monitoring the cell death macroscopically (Figure 2D) and microscopically using dead-cell-staining trypan blue at 20 hpi (Figure 2E), the infiltration sites of *H. chejuensis* and *P. syringae* pv. tomato did not differ. *PR-1a* gene expression in *N. benthamiana* (*NbPR-1a*) for *H. chejuensis* and *P. syringae* pv. tomato implicated that the HR-like cell death was a result of active defense (Figure 2F).

### HR Suppressed by *H. chejuensis avrPto1* and *NbSGT1* Silencing

AvrPto1, a T3SS-secreted effector of *P. syringae* pv. tomato, inhibits HR induced by nonhost-pathogen interactions (Kang et al., 2004). Thus, we examined whether by *H. chejuensis*-induced HR-like cell death in *N. benthamiana* would be suppressed by *H. chejuensis* expressing *avrPto1*. As expected, infiltration of *H. chejuensis* harboring *avrPto1* suppressed HR-like cell death (Figure 2G), indicating *avrPto1* expression in *H. chejuensis* and AvrPto1 translocaion into the cytoplasm of *N. benthamiana* through T3SS. To verify that the HR-like cell death follows a typical defense-associated PCD mechanism, we examined the cell death after virus-induced gene silencing of *N. benthamiana SGT1* (*NbSGT1*). SGT1 is a general regulator of plant resistance, and silencing *NbSGT1* compromises HR development during R-gene-mediated and nonhost resistance (Peart et al., 2002). The cell death caused by *H. chejuensis* was completely abolished in *NbSGT1*-silenced *N. benthamiana* (Figure 2G). These observations demonstrate that the PCD of *N. benthamiana* caused by *H. chejuensis* is a result of nonhost resistance.

## DISCUSSION

In this study, we provide the first evidence that a land plant, *N. benthamiana*, can respond to the marine bacterium *H. chejuensis* with PCD. Based on the presence of animal-pathogen-type T3SSs and other genes that may play roles in virulence (Jeong et al., 2005), we speculate that *H. chejuensis* may be a pathogen or a symbiont of a marine protist or animal. Because of their different habitats, it is unlikely that *H. chejuensis* and *N. benthamiana* encounter under natural conditions. Nevertheless, *N. benthamiana* exhibited AvrPto1-inhibited SGT1-mediated PCD against *H. chejuensis*. Thus, we hypothesize that *N. benthamiana* responded either directly to the effectors translocated through T3SS or more likely to the protein modification or metabolic perturbation caused by the activities of translocated effectors, which are also recognized by NLR proteins. It is improbable if not impossible that *Hahella* effectors absent in plant-pathogenic bacteria or phylogenetically distant are specifically recognized by the plant NLR protein. NLR-triggered defense response is often dependent on recognition of effector-mediated disturbances of target proteins or homeostasis as described in the guard and decoy hypothesis (Mukhtar et al., 2016). A recent study suggests that certain NLRs form complex networks for sensing pathogen invasion and eliciting plant immune responses (Duxbury et al., 2021), and the networks may have functionally redundant nodes that can be targeted by pathogen effectors (Adachi et al., 2019; Derevnina et al., 2021).

To our knowledge, this is the first study to document that a T3SS-harboring marine bacterium elicits PCD in a land plant, which is an extreme example of nonhost resistance. Our work that provides novel insights into the inner workings of plant immunity sheds lights on the evolution of plant defense systems. It remains to be investigated how the plant has acquired the trait to recognize a potential threat posed by unprecedented T3SS effectors, and which receptor(s) is/are responsible for the recognition through what mechanism.

## MATERIALS AND METHODS

### Plants

*Nicotiana benthamiana* was grown in a plant growth chamber (Sejong Biotech. Korea) at 25 °C under a photoperiod of 16 h. When they reached the six-leaf stage, the plants were subjected for HR-like response assays.

### Bacterial cultures and infiltration

*Hahella chejuensis* was grown in LB or LB containing 3% NaCl medium supplemented with appropriate antibiotics at 30 °C. *Escherichia coli* EPI300 was grown in LB medium (BD Difco™) at 37 °C, and *P. syringae* pv. *tomato* DC3000 T1a was grown on King’s B medium (10% proteose peptone, 1.5% K_2_HPO_4_, 15% glycerol, and 5mM MgSO_4_) or in LB broth at 30 °C. *Agrobacterium* GV3101 containing TRV-VIGS vectors were grown in LB at 28 °C. To identify the HR-like response elicited by *H. chejuensis* on *N. benthamiana* at different stages of growth, fresh cultures were diluted in LB medium containing 3% NaCl and grown at 30 °C until the cultures reached the stationary phase. Bacteria samples were taken for inoculation of *N. benthamiana* at every 2 h time intervals. Each sample was centrifuged and resuspended in PBS at a cell density of 1 × 10^8^ CFU/mL and infiltrated into *N. benthamiana* leaves. Infiltrations were performed by pricking leaves with a needle and then pressing the needless syringe against the leaf surface while supporting the leaf with a finger (Chaudhry et al., 1987). The development of a HR-like response was observed after 40 h at room temperature.

### Construction of a virulence gene tree

The amino acid sequences of prokaryotic type-III secretion inner membrane channel proteins (StcV, EscV, LcrD, HrcV, and SsaV) were downloaded from the UniProtKB database. In total, 509 protein sequences were aligned using MUSCLE software (Edgar, 2004) with the complete gap deletion option, and evolutionary analyses were conducted using MEGA X (version 10.1.5) (Kumar et al., 2018) based on the Maximum Likelihood method and a JTT matrix-based model. In total, 381 amino acid positions were used to infer evolutionary distances. The constructed phylogenetic tree was exported in Newick tree format and was visualized using iTOL (http://itol.embl.de) (Letunic and Bork, 2019).

### Plasmids

The plasmid pPTE6::*AvrPto1* was transformed into *H. chejuensis* by electroporation and was selected on LB medium containing kanamycin (50 ng/mL) at 30 °C. *H. chejuensis* (*AvrPto1*) transformants were confirmed through PCR using specific *AvrPto1* primers in supplemental table S1.

### Quantitative RT-PCR of T3SS RNA in *H. chejuensis*

For quantitative reverse transcription PCR, Total RNA was extracted from *H. chejuensis* using the RNeasy plus Mini Kit (Qiagen, Germany) and was eluted in RNase-free water. Reverse transcription was carried out using 400 ng total RNA followed by quantitative PCR using iQ™ SYBR Green Supermix (BioRad, USA). To calculate the relative fold change, the 2^-ΔΔCT^ method was employed using the threshold cycle (CT) values obtained from qPCR of each T3SS genes at different time points, CT value of 4h sample which is the earliest and housekeeping gene *rpoA* in *H. chejuensis*. All primers used are listed in Supplemental Table S1.

### Microscopic observations and trypan blue staining

The leaf disks containing inoculum were excised at 12 h after inoculation and stained with lactophenol-trypan blue (10 mL of lactic acid, 10 mL of glycerol, 10 g of phenol, and 10 mg of trypan blue dissolved in 10 mL of distilled water (Keogh et al., 1980)). Whole leaves were boiled for 1 min in the staining solution and then decolorized in chloral hydrate (2.5 g of chloral hydrate dissolved in 1 mL of distilled water) for at least 30 min. The bleached (stained) leaves were examined by microscopy under a compound microscope equipped with interference or phase-contrast optics.

### *PR-1a* Gene Expression in *N. benthamiana*

Total RNA was extracted from the leaves of *N. benthamiana* 12 h after inoculation with *H. chejuensis* and *P. syringae* pv. *tomato* DC3000 as a positive control and PBS as a negative control. For total RNA isolation in plant tissue, leaf disks were collected with a 1 cm diameter cork borer, frozen in liquid nitrogen, ground into powder, and extracted RNA using Trizol reagent (Invitrogen) and suspended in water. Reverse transcriptase reactions were carried out using 1 µg of total RNA. We used 1 µl of the reverse transcription reaction as a template in 20 µl PCR reactions with primer sets listed in Supplemental Table S1. The RT−PCR products were separated by agarose gel electrophoresis.

### Virus Induced Gene Silencing (VIGS) through *NbSGT1*

*Agrobacterium* GV3101 containing TRV-VIGS vectors either without an insert (TRV::00) or with an *N. benthamiana SGT1* insert (TRV::*NbSGT*) were grown overnight, and cultures were harvested by centrifugation at 4,000 rpm, followed by washing once in 10 mM MES (pH 5.6), and resuspension in *Agrobacterium* inoculation buffer (10 mM MgCl_2_, 10 mM MES (pH 5.6), and 200 µM acetosyringone) to a final concentration of 1 × 10^8^ CFU/mL. These mixtures were incubated for 4 h under shaking at room temperature before infiltration. *Agrobacterium* cultures containing TRV-VIGS vectors were infiltrated into the leaves of 14-day-old tobacco plants using 1 mL needleless syringes. After two to three weeks of VIGS, the plants were subjected to *H. chejuensis-* treatments.

## SUPPLEMENTAL DATA

Supplemental Table S1. Primer sequences used in this study

## ACKNOWLEDGMENTS

We thank S. P. Dinesh-Kumar for providing GATEWAY ready TRV-VIGS vectors, Alan Collmer for pCPP3221 plasmid. This work was supported by grants from the National Research Foundation of Korea (2018R1A6A1A03025607 and RS-2023-00211512) and the KRIBB Research Initiative Program.

## AUTHOR CONTRIBUTIONS

J.F.K., C.-M.R., and S.K.K. conceived, organized, and supervised the study. B.L., J.-I.L., and S.K.K. performed the experiments. S.K.K. conducted sequence comparison and phylogenetic study. B.L., S.K.K., C.-M.R., and J.F.K. wrote and edited the manuscript.

